# Phased ERK function on muscle stem cell plasticity in axolotl

**DOI:** 10.1101/2025.04.30.650958

**Authors:** Chao Yi, Xinxin Liu, Liqun Wang, Li Song, Ji-Feng Fei

## Abstract

The precise regulation of muscle stem cell (MuSC) plasticity remains poorly understood. While studies have suggested that *Pax7*+ MuSCs in vertebrate limbs are committed solely to myogenic lineages during regeneration, we previously observed during axolotl tail regeneration, *Pax7*+ MuSCs contribute to the fibroblast and chondrocyte mesodermal lineages. However, the underlying mechanism is unclear. Here, we show that the duration of ERK signaling plays an important role in MuSCs cell fate. We observed that ERK exhibits sustained activation after tail amputation that is followed by subsequent inhibition. We found that ERK activation initiates MuSCs plasticity switch toward a trunk fibroblast fate by repressing PAX7 expression. However, subsequent inhibition of ERK activity is essential for the transition of trunk fibroblast to fin fibroblast. In addition, we found that the TGF-β/SMAD2 cascade as a downstream mediator of ERK-driven regulation of MuSC plasticity towards fibroblast and chondrocyte during tail regeneration. Together, we uncovered a new phased stem cell regulatory mechanism linking injury-induced ERK signaling to stem cell plasticity regulation, offering new insights with potential implications for regenerative medicine.

## INTRODUCTION

Precise regulation of stem cell plasticity is critical for tissue homeostasis and regeneration, as insufficient plasticity can hinder repair while excessive plasticity may predispose to tumorigenesis.^1,2,3^ In species with limited regenerative capacity, such as mice, adult muscle stem cells (MuSCs) are predominantly committed to myogenesis for homeostasis and repair.^4,5,6^ Under some conditions, for instance, *Pax7*/*Pax3* knockout, aging, or in vitro cell culture expansion, MuSCs can be reprogrammed to adopt non-myogenic fates, such as chondrocytes, fibroblasts, mesenchymal cells, or adipocytes.^7,8,9,10^

In highly regenerative species such as newts, transplantation studies demonstrate that MuSCs can differentiate into non-myogenic lineages following their isolation, in vitro expansion, and reimplantation into the regenerating limb blastema.^11,12^ However, the process of in vitro culture itself may predispose satellite cells towards mesenchymal plasticity.^10^ In axolotl, we have recently provided the first direct in vivo lineage-tracing evidence that the plasticity of *Pax7*+ MuSCs differs between homeostasis and regeneration in the tail. Specifically, while tail MuSCs remain restricted to the muscle lineage during homeostasis, they acquire the capacity to differentiate into fibroblasts, chondrocytes, and pericytes during regeneration.^13^ However, the mechanisms that stimulate MuSCs to acquire alternative differentiation fates remain unknown.

The regulation of MuSC lineage commitment towards the muscle fate involves multiple modulators, including transcription factors,^14,15,16^ microenvironment stiffness,^17^ autophagy, secreted factors,^18^ metabolites, and epigenetic modifications.^19,20^ In contrast, although non-myogenic plasticity of MuSCs has been observed in vitro, and aging-induced WNT signaling has been implicated in driving fibrogenic conversion in mice,^8^ direct in vivo evidence delineating the signaling mechanisms that govern MuSC plasticity toward fibroblast differentiation is still lacking.

Here, we demonstrate that in axolotl tail regeneration the transition of *Pax7*+ MuSCs plasticity from myogenesis to fibrogenesis is orchestrated by phased ERK activity. During the early regenerative stage, sustained ERK activation inhibits PAX7 and is sufficient to induce MuSCs to generate trunk fibroblasts. Conversely, ERK inhibition during the fin outgrowth stage promotes the conversion of trunk fibroblasts to fin fibroblasts. Moreover, we also demonstrate that during regeneration, ERK modulates trunk fibroblast differentiation through SMAD2, but not SMAD3. Collectively, these results identify ERK as a molecular switch whose dynamic regulation is essential for altering the original monopotent muscle differentiation fate of MuSCs to a differentiation state to fibroblasts in regenerating tails of axolotls.

## RESULTS

### ERK activity and PAX7 expression exhibit opposing temporal profiles in the *Pax7*+ MuSC lineage during axolotl tail regeneration

To assess whether muscle stem cell (MuSC) plasticity is altered during axolotl tail regeneration, we lineage-traced *Pax7*+ MuSCs using a previously established transgenic model (*Pax7:CreERT2* x *CAGGS:reporter*) during both tail development and regeneration (Figure 1A).^21,22^ After a two-week tamoxifen (4-OHT) induction, we analyzed the progeny of *Pax7*+ MuSCs at 21 days post-treatment (dpt) during development, as well as at 6 days post-amputation (dpa) representing the early regenerative stage and 15 dpa representing the fin outgrowth late stage (Figure 1B). During development, MuSCs contributed exclusively to muscle formation (Figure 1Ca), which is also confirmed by our previous work.^13^ In contrast, during regeneration, MuSCs first generated trunk fibroblasts at 6 dpa (Figure 1Cb, Figure S1A) and later produced fin fibroblasts and muscle cells at 15 dpa (Figure 1Cc, Figure S1B). Collectively, these observations indicate that MuSC differentiation potential shifts from a solely myogenic fate during development to extra fibrogenic lineage during tail regeneration.

**Figure 1.**
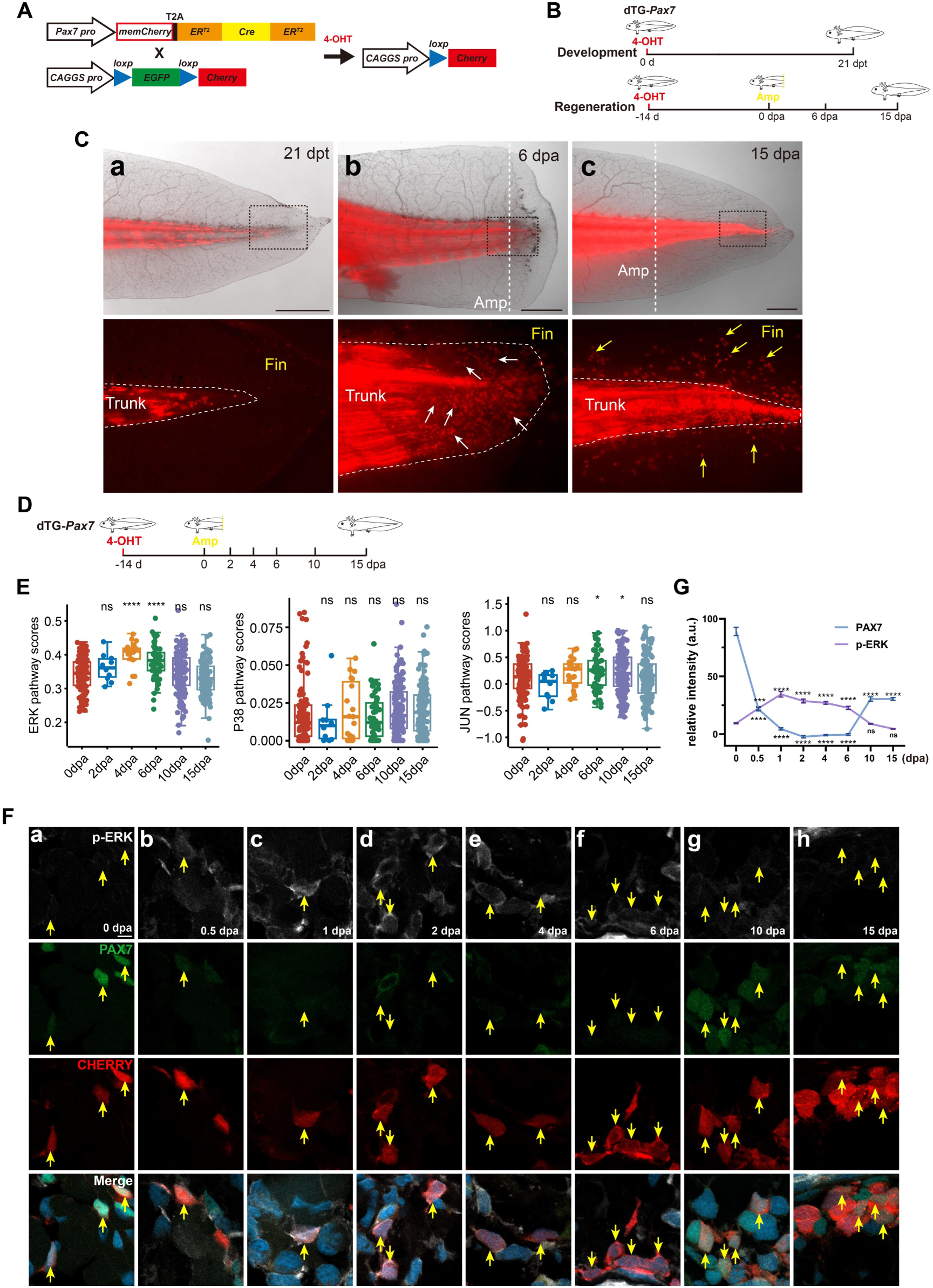
Opposing dynamics of ERK activity and PAX7 expression in the *Pax7*+ muscle stem cell (MuSC) lineage during axolotl tail regeneration. (A, B) Schematic of the experimental strategy for lineage tracing *Pax7*+ MuSCs during development and regeneration. Amp: amputation; dpt: days post 4-OHT treatment; dpa: days post amputation. (C) During development, *Pax7*+ MuSCs exclusively generated muscle (a), whereas during regeneration they differentiated first into trunk fibroblasts (b, white arrows) and then fin fibroblasts (c, yellow arrows) (n > 6 per group). The rectangles were shown at higher magnification below. Dashed lines marked the amputation sites in (b, c). (D) Schematic of single-cell RNA sequencing performed on lineage-traced *Pax7*+ MuSCs and their progeny at the indicated time points. (E) Among MAPK pathways, only ERK displayed marked fluctuations during regeneration, in contrast to p38 MAPK and JNK. (F) Immunostaining revealed dynamic changes in ERK activity and PAX7 expression within the MuSC lineage during tail regeneration. (G) Quantification of pERK and PAX7 signals in (F) (mean ± SEM; n=3; ***p < 0.001; ****p < 0.0001; ns, not significant). Scale bars: 2 mm (C); 10 μm (F).

To elucidate the signaling mechanisms governing this plasticity switch, we looked at up-regulated signals within MuSCs lineage during axolotl tail regeneration. From our published literature, WNT, MAPK, and TGF-β were among the top three significantly altered pathways during axolotl tail regeneration.^13^ Notably, the MAPK pathway emerged as the most pertinent to injury response due to its rapid activation and direct integration of damage signals.^23,24^ To further dissect MAPK subfamily activation, we re-analyzed our publicly available single-cell RNA-seq dataset of the *Pax7*+ MuSC lineage sampled at defined stages of tail regeneration (Figure 1D).^13^ Our results demonstrated that ERK, not JNK or p38 MAPK, is predominantly activated during the early regenerative stages (Figure 1E).

To confirm ERK dynamics in MuSCs lineage during axolotl tail regeneration, we monitored ERK activity in *Pax7*+ MuSCs and their progeny using the *Pax7:CreERT2* x *CAGGS:reporter* animals. Co-immunostaining for phosphorylated ERK (p-ERK) and the MuSC marker PAX7 revealed that ERK activity was markedly elevated in MuSCs during the early immune response and wound healing phase (0.5-1 dpa) (Figures 1Fa-1Fc). During early regeneration stage (2-6 dpa), ERK activation remained sustained in MuSC-derived progeny (Figures 1Fd-1Ff), whereas in th fin outgrowth stage (10-15 dpa), ERK activity was significantly reduced to a level similar to uninjured controls (0 dpa) (Figures 1Fg-1Fh). In contrast, PAX7 expression exhibited an inverse pattern to that of pERK (Figures 1F and 1G). Moreover, analysis of MuSCs within a 500 µm regeneration-responsive zone (Figure S2A)^25,26^ proximal to the amputation site at 0.5 dpa demonstrated that ERK activity was substantially higher near the injury compared to distal regions, while PAX7 levels were lower, suggesting a localized activation gradient (Figures S2B-S2D). Collectively, these findings show that ERK activity in the MuSCs lineage is initially sustained and later inhibited during tail regeneration, indicating a time-regulated function in the progeny of *Pax7*+ MuSCs during the regenerative process.

### ERK activation initiates MuSCs plasticity towards trunk fibroblasts during homeostasis

To test the role of sustained ERK activation on MuSC plasticity in axolotl tail, we constructed constitutively active (caMEK1) and dominant negative (dnMEK1) mutants of axolotl MEK1 (the upstream kinase of ERK) to precisely modulate ERK signaling (Figure S3A). Based on cross-species conservation (Figure S3B), we generated MEK1 mutants in which Ser218 and/or Ser222 were replaced with aspartate (to mimic phosphorylation) or alanine (to mimic dephosphorylation).^27,28,29^ In cell culture, caMEK1(S218, 222D) strongly promoted the phosphorylation of endogenous ERK (Figure S3Ca and S3Cd), while dnMEK1 (S218, 222A) effectively inhibited it (Figure S3Cc). We observed that the inhibitory effect of dnMEK1 (S218,222A) was stronger than dnMEK1 (S222A) (Figures S3Cb and S3D). According to this, we took caMEK1(S218, 222D) and dnMEK1 (S218,222A) and generated two double transgenic lines (*Pax7:CreERT2/CAGGS:loxp-STOP-loxp-caMek1-Cherry* and *Pax7:CreERT2/CAGGS:loxp-STOP-loxp-dnMek1-Cherry*) that will allow us to conditionally modulate ERK activity in MuSCs with 4-OHT treatments (Figure 2A). We then activated or inhibited ERK in MuSCs for 21 days, and analyzed the fate of MuSCs fate during homeostasis (Figure 2B). After 21 days of sustained ERK activation, we observed that *Pax7*+ MuSCs gave rise mostly to trunk fibroblasts cells (Figures 2Ce and 2Cf), which were located between dermis and muscle compartment, with expression of PRRX1 (Figure 2Dc), a marker for fibroblast generally used in axolotls.^30,31^ After sustaining ERK activation for 30 days, we even observed more truck fibroblast formation, replaced the original myotome (Figure S4Ac). In contrast, in the control and ERK inhibition groups, we observed only the contribution of MuSCs to muscle cells (Figures 2Ca-2Cd; Figures 2Da and 2Db, Figure S4Aa and S4Ab). By Tracing the same population of *Pax*7+ MuSCs lineage over time revealed that, after ERK activation, the cells that harbored fibroblast morphology were most restricted to the trunk: only a few cells migrated into the fin (Figure S5C). In the control and ERK inhibition groups, however, cells remained confined to the muscle compartment, maintaining muscle cell morphology (Figure S5A, S5B).

**Figure 2.**
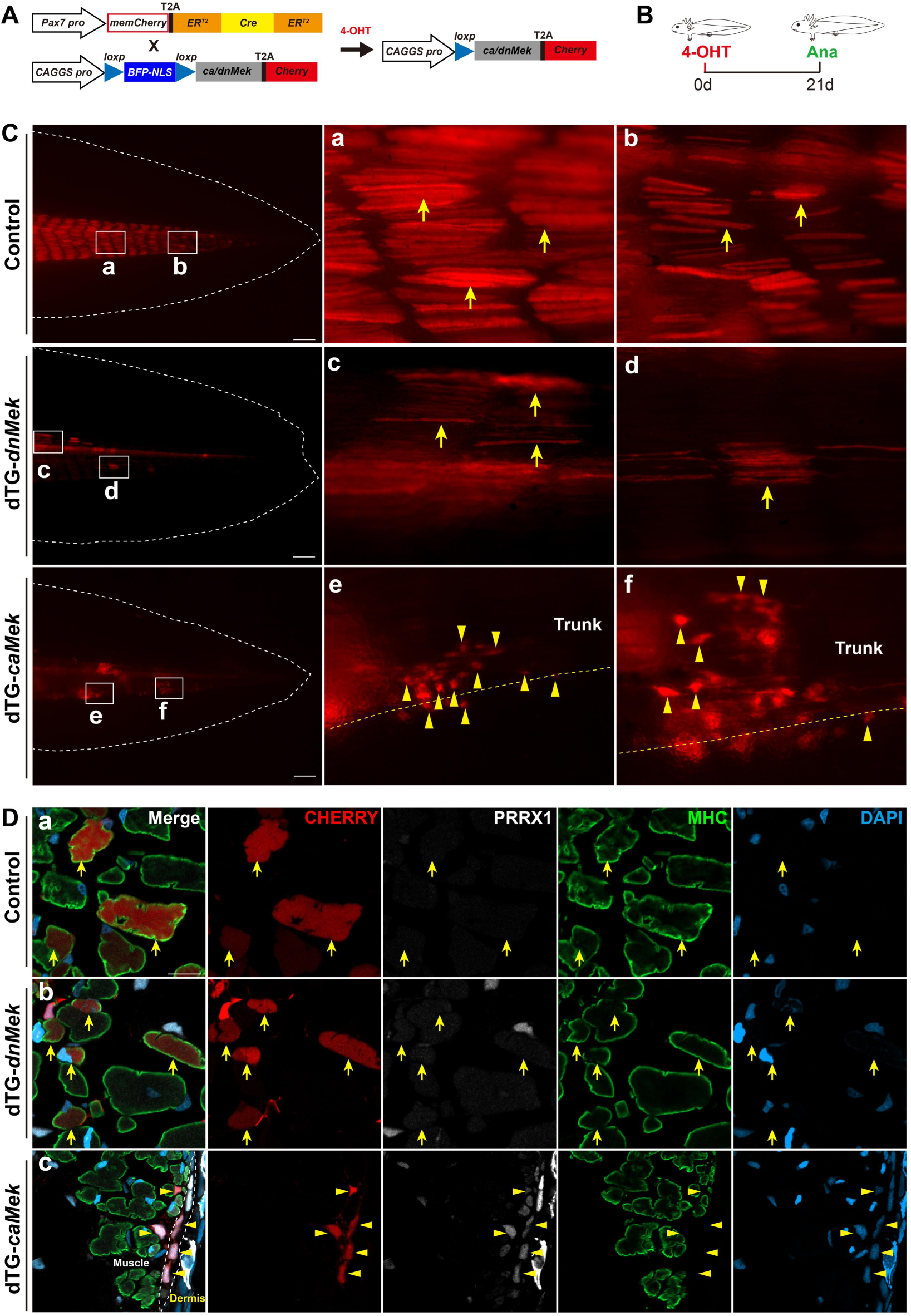
Sustained ERK activation drives *Pax7*+ MuSC plasticity towards a trunk fibroblast fate. (A, B) Experimental strategy for manipulating ERK signaling in the MuSC lineage. (C) Representative images demonstrating that, in the absence of injury, enforced ERK activation (dTG-*caMek*, n=10) promoted the generation of trunk fibroblasts, whereas ERK inhibition (dTG-*dnMek*, n=7) and Cherry controls (n=6) resulted exclusively in muscle formation. Dashed white and yellow lines delineated the fin shape and the trunk region. The rectangles were shown at higher magnification at the right side. (D) Molecular characterization of MuSC progeny using immunostaining for PRRX1 (fibroblast marker) and MHC (muscle marker). Dashed lines in (c) indicated the dermal trunk fibroblasts region. Scale bars: 100 μ m (C); 25 μ m (D). Yellow arrows denote muscle cells; yellow triangles denote trunk fibroblasts.

To determine whether ERK activation regulates MuSC differentiation towards the fibroblast lineage in a conserved manner, we evaluated MuSCs fate in the axolotl head and limb following 30 days of ERK activation. Similarly, we observed fibroblast cells derived from MuSCs in both the head (Figure S4Af) and limb (Figure S4Bf). Further immunostaining on limb MuSCs derived progeny confirmed their identity as PRRX1+ fibroblasts, with no detectable MHC or MEF2C muscle-lineage markers (Figure S4C). Surprisingly, we also detected SOX9+ chondrocytes from MuSCs-derived progeny in the limb after ERK activation (Figure S4Bc). To further explore the conservation of this mechanism, we tested the effect of ERK activation in C2C12 mouse myoblast cells. We established a stable C2C12 cell line expressing caMEK and confirmed increased ERK activation by p-ERK immunostaining (Figure S6A). After 4 days of ERK activation and induction of differentiation with low serum, we observed a significant reduction in muscle cells length, accompanied by a marked decrease in the muscle marker MHC (Figure S6B-S6D). These results suggest that ERK-mediated regulation of MuSC plasticity is conserved across different tissues and species.

Furthermore, to test whether WNT or TGF-β pathway initiate MuSC differentiation towards fibroblasts, we constructed two Cre-LoxP lines: *Pax7:CreERT2/CAGGS:loxp-STOP-loxp-caCtnnb1-Cherry* to activate WNT signaling, and *Pax7:CreERT2/CAGGS:loxp-STOP-loxp-caTgfbr1-Cherry* to conditionally activate TGF-β in *Pax7*+ MuSCs (Figures S7A and S7B; Figures S7D and S7E).^32^ However, at 21 days post-4-OHT treatment, we did not observe fibroblasts derived from MuSCs in either the WNT or TGF-β activation groups (Figure S7C; Figure S7F).

In summary, these findings suggest that sustained ERK activation is a conserved mechanism regulating MuSCs plasticity. The activation of ERK, but not WNT or TGF-β, is sufficient to reprogram *Pax7*+ MuSCs plasticity towards trunk fibroblasts, a response observed specifically during early stages of fin regeneration (6 dpa) (Figure 1Cb).

### ERK activation inhibits PAX7 expression, triggering MuSCs Plasticity towards trunk fibroblasts during tail regeneration

We observed that ERK activity and PAX7 expression were mutually exclusive during axolotl tail regeneration: PAX7 levels decreased as ERK activity increased, and reappeared when ERK activity declined (Figure 1G). This inverse correlation suggests that ERK may negatively regulate PAX7 expression, a relationship that has been reported with conflicting outcomes in mammalian studies.^33,34,35^ To address this in axolotls, we conditionally manipulated ERK activity in MuSCs (Figure S8A). At 7 and 14 days post-4-OHT treatment, ERK activation significantly reduced PAX7 expression in MuSCs, whereas ERK inhibition did not alter PAX7 levels (Figure S8B-S8G). Upon tail amputation, PAX7 expression soon decreased (Figure 1F, 1G). However, ERK inhibition upregulated *Pax7* expression comparing to control animals at 6 hours post tail amputation (Figure S8H, 8I). Together, these findings indicate that ERK activation suppresses PAX7 expression in axolotl MuSCs.

To test whether loss of PAX7 alone can mimic the ERK-dependent shift of MuSC plasticity towards a fibrogenic fate, we assessed MuSCs fate following knockout of endogenous PAX7. We took advantage of the previously established *Pax7:Cherry* knock-in/knockout axolotl model, which enables identification and fate tracing of PAX7 knockout MuSCs via the Cherry reporter.^36^ Compared to PAX7+/+ animals, PAX7+/- animals showed a clear reduction in PAX7 levels and an increasement in PRRX1 expression (Figure 3A, 3B, 3D, 3E). In PAX7+/+ animals, only 2 of 121 traced cells (1.6%) exhibited PRRX1 intensity equivalent to trunk fibroblasts. Heterozygous loss of PAX7 increased this proportion modestly (8/85 cells; 9.4%). Strikingly, complete ablation of PAX7 (PAX7−/−) drove robust PRRX1 upregulation in Cherry+ MuSCs and their progeny, with 65 of 133 cells (48.9%) matching trunk fibroblast PRRX1 levels and localizing predominantly within the trunk region (Figure 3C-3E). These results demonstrate that the lack of PAX7 is sufficient to induce MuSCs plasticity toward a trunk fibroblast fate.

**Figure 3.**
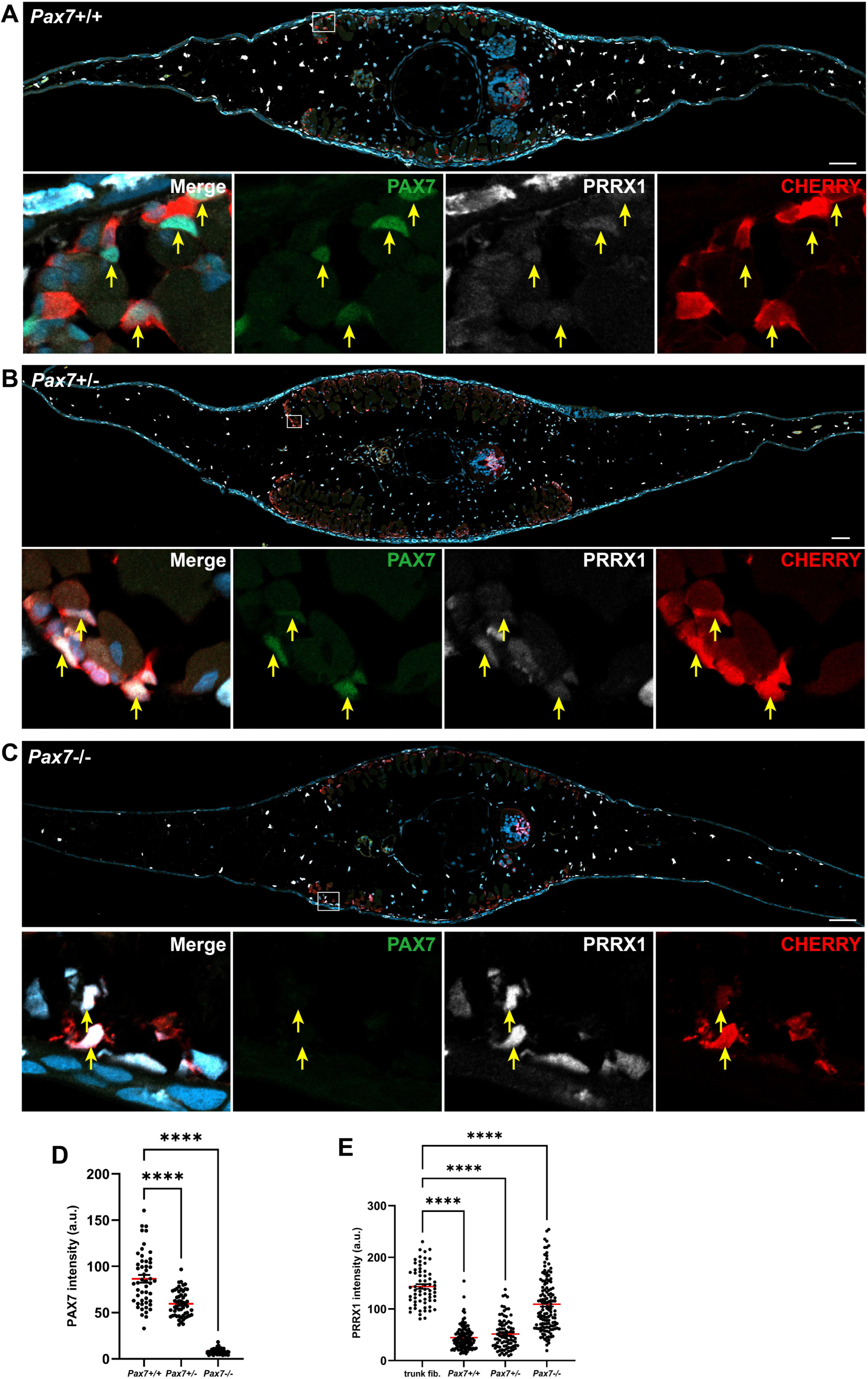
PAX7 knockout directs MuSC fate towards a trunk fibroblast lineage. (A-C) Representative images displayed PAX7 and PRRX1 expression in lineage-traced wild type animals (A, n=6), PAX7 knockout heterozygotes (B, n=6), and PAX7 knockout homozygotes (C, n=6). Arrows indicate MuSCs; inset panels show enlarged views of the boxed areas. (D-E) Quantitative analysis of PAX7 (D) and PRRX1 (E) expression in MuSCs (mean ± SEM; each dot represents a single cell; ****p < 0.0001). Scale bars: 100 μm.

### Inhibition of ERK activity is required for trunk fibroblasts transition to fin fibroblasts

Axolotl tail regeneration is characterized by the progressive conversion of trunk fibroblasts into fin fibroblasts from early to late regenerative stages (Figures 1Cb-1Cc). Notably, fin fibroblasts express higher levels of PRRX1 compared to trunk fibroblasts (Figures 4A and 4B), and ERK activity exhibits a corresponding phased pattern: initially activated and subsequently inhibited during regeneration (Figure 1G). In contrast, sustained ERK activation in MuSCs and their progeny inhibited the formation of fin fibroblasts (Figures 2Ce and 2Cf; Figures S4Ac, S5C) and we observed suppression in PRRX1 expression (Figures 4C and 4D). These observations led us to hypothesize that ERK inhibition is essential for the conversion of trunk fibroblasts into fin fibroblasts by regulating PRRX1.

**Figure 4.**
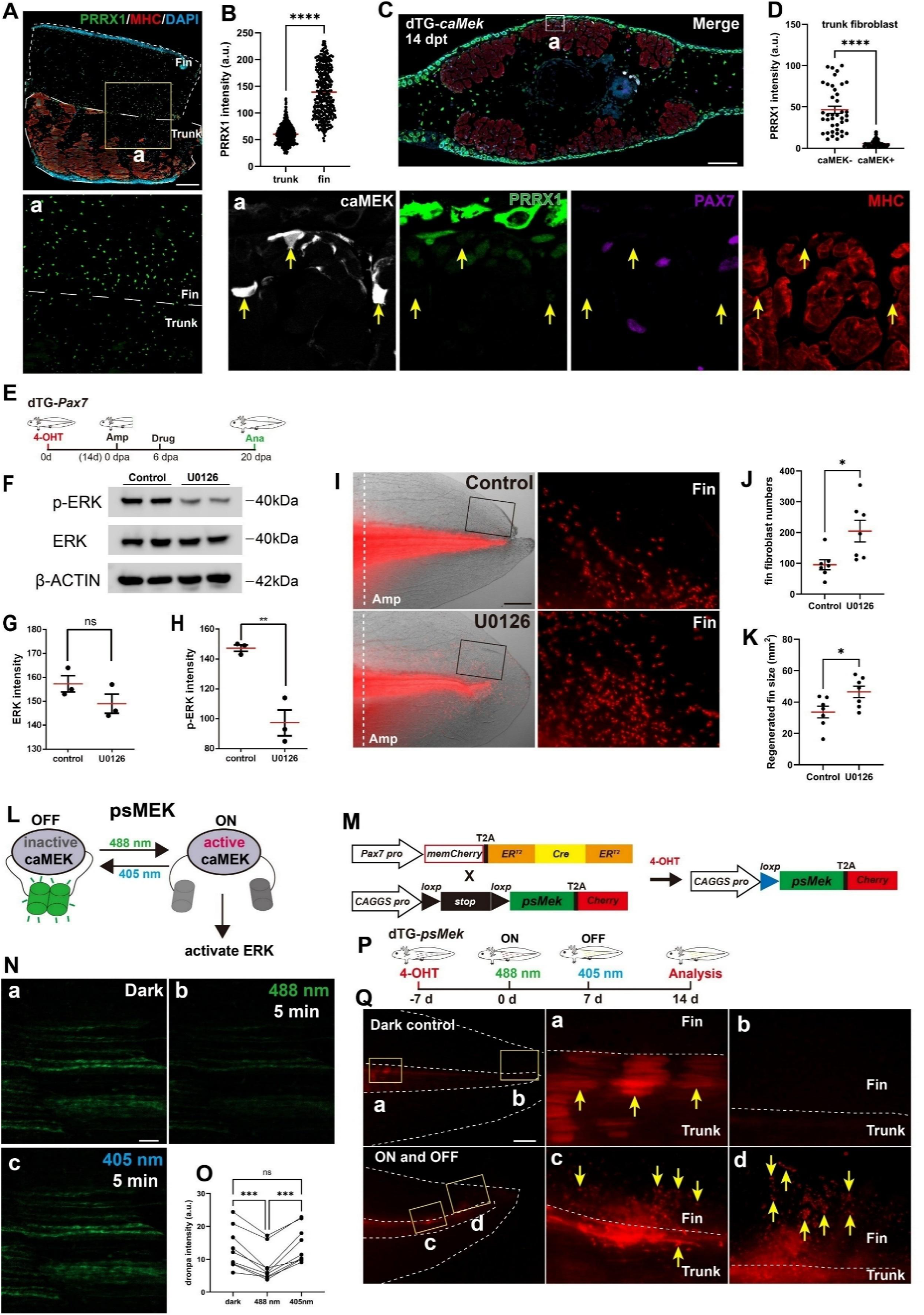
ERK inhibition is required for fin fibroblast generation. (A) Immunostaining for fibroblasts (PRRX1) and muscle (MHC) in uninjured wild-type axolotls; dotted lines delineate trunk and fin regions (n=6). Enlarged image was shown in (a). (B) Quantitative analysis of PRRX1 intensity from (A) (mean ± SEM; each dot represents a single cell; ****p < 0.0001). (C) Representative images at 14 dpt showing that caMEK-expressing cells (arrows in Ca) exhibit reduced PRRX1 expression (n=7). Enlarged images were shown in (a). (D) Quantitative comparison of PRRX1 intensity between caMEK negative (caMEK-) and caMEK positive cells (caMEK+) in (C) (mean ± SEM; each dot represents a single cell; ****p < 0.0001). (E) Schematic of the experimental strategy for pharmacologically inhibiting ERK activity after trunk fibroblast emergence. (F) Western blot analysis of pERK and total ERK under U0126 (ERK inhibition) and DMSO (control) treatment conditions. (G, H) Quantitative analysis of pERK and ERK levels from (F) (mean ± SEM; each dot represents an experiment; **p < 0.01; ns, not significant). (I) Representative images of fin fibroblast distribution (Cherry-positive cells) at 20 dpa in control and ERK inhibition groups; white dashed lines indicate amputation sites, and insets show enlarged views of the boxed regions. (J, K) Quantitative analysis of fin fibroblast numbers and regenerated fin size from (I) (mean ± SEM; each dot represents an animal; *p < 0.05). (L) Schematic describing the optogenetic tool psMEK. (M) Schematic of the strategy for generating a transgenic line to optogenetically modulate ERK activity in the MuSC lineage while tracing cell fate. (N) Confocal live imaging demonstrating effective on/off switching of psMEK under 488 nm or 405 nm illumination for 5 minutes. (O) Quantitative analysis of Dronpa intensity from (N) (mean ± SEM; each dot represents an animal; ***p < 0.001; ns, not significant). (P) Schematic outlining the optogenetic strategy for manipulating ERK activity. (Q) Representative images showing fin fibroblast generation following sequential ERK activation and inhibition (n=4) compared to dark controls (n=3); Arrows indicate muscle in (a) and fin fibroblasts in (c, d). Scale bars: 500 μm (A); 200 μm (C); 1 mm (I, Q); 100 μm (N).

To test this hypothesis, we first inhibited endogenous ERK activity using drug U0126. In *Pax7:CreERT2* x *CAGGS:reporter* transgenic axolotls—where *Pax7*+ MuSCs were labeled at 14 days post-4-OHT treatment—we administered U0126 from 6 to 20 days post-amputation, a window during which trunk fibroblasts had formed but fin fibroblasts had not yet emerged (Figure 4E). At 20 dpa, ERK inhibition by U0126 was confirmed by western blotting (Figures 4F-4H), and we observed an increase in fin fibroblast numbers compared to controls, accompanied by a significant increase in fin size—presumably due to the enhanced fibroblast population (Figures 4I-4K). These data support the role of ERK inhibition in promoting the transition from trunk to fin fibroblasts.

To rule out off-target effects of U0126 and further pinpoint the role of ERK in MuSCs, we employed an optogenetic tool, psMEK (photoswitchable MEK), which allows precise temporal control of ERK activity.^37^ The psMEK construct incorporates two light-switchable Dronpa domains fused to caMEK. Under 405 nm illumination, Dronpa dimerizes and inactivates caMEK, whereas 488 nm illumination dissociates the dimer, reactivating caMEK and thereby endogenous ERK (Figure 4L). We generated *Pax7:CreERT2/CAGGS:loxp-STOP-loxp-psMek-Cherry* transgenic animals so that, following 4-OHT induction, we could toggle ERK activity in MuSCs and trace their fate via the Cherry reporter (Figure 4M). In these animals, 5 minutes of 488 nm illumination dissociated the Dronpa dimer and activated ERK (Figures 4Nb, S9Aa), while 408 nm illumination re-induced dimerization to suppress ERK activity (Figures 4Nc, 4O; Figures S9Ab and S9B). We then maintained ERK activity for 7 days followed by 7 days of inhibition to mimic the observed endogenous changes in ERK activity during regeneration (Figure 4P). Compared to dark controls, which produced only muscle cells (Figures 4Qa and 4Qb, S9Ca), animals subjected to this light regimen generated fin fibroblasts with high PRRX1 expression (Figures 4Qc and 4Qd, S9Cb and S9D).

Collectively, these results demonstrate that inhibition of ERK activity is a critical trigger for the transition of trunk fibroblasts to fin fibroblasts during axolotl tail regeneration.

### ERK regulates MuSC plasticity via TGF-β signaling during axolotl fin regeneration

To investigate the role of ERK activity in directing MuSC fate during regeneration, we manipulated ERK signaling in PAX7+ MuSCs and their progeny and traced their lineage throughout tail regeneration (Figure 5A). Consistent with our previous observation,^13^ axolotl MuSCs give rise to both muscle cells, fibroblasts and chondrocytes by 21 dpa (Figures 5Ba, 5Ca and 5Cb). In contrast, dnMEK-Cherry recombined animals produced exclusively muscle cells (Figures 5Bb, 5Cc), while caMEK-Cherry recombined axolotls exhibited a shift toward fibroblast and chondrocyte differentiation (Figures 5Bc 5Cd and 5Ce), Notably, consistent with what we observed above, PRRX1 expression level is inhibited (Figure 5Cd). These results indicate that ERK signaling functions as a molecular switch that determines whether MuSC differentiate into muscles or fibroblasts.

**Figure 5.**
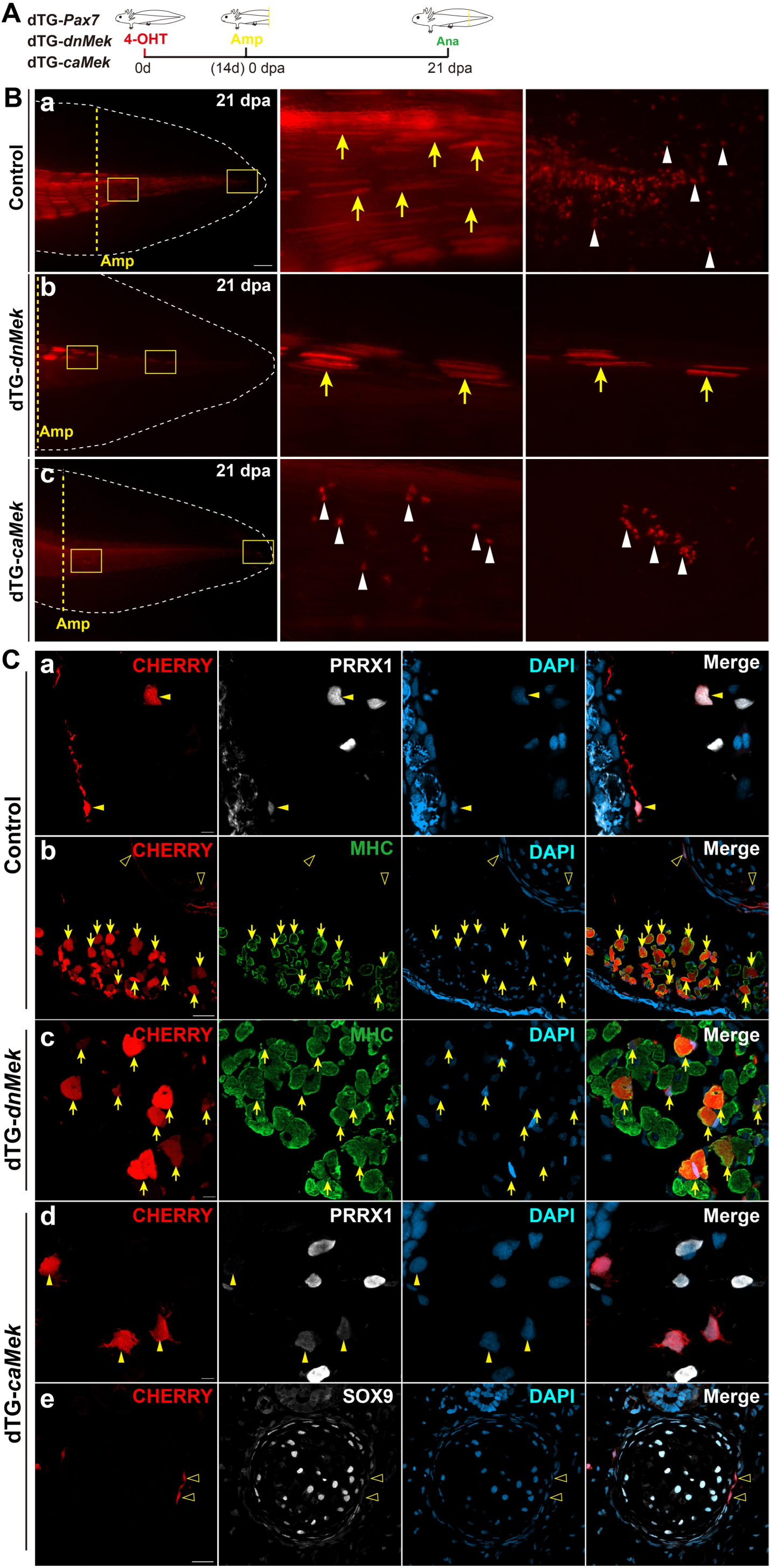
ERK directs MuSC fate decisions between muscle and fibroblasts during axolotl tail regeneration. (A) Schematic of the experimental strategy for modulating ERK signaling during tail regeneration. (B) Representative images of MuSC-derived cells in control (a) (n=6), continuously ERK-inhibited (b) (dTG-*dnMek,* n=3), and continuously ERK-activated (c) (dTG-*caMek*, n=4) animals at 21 dpa. Yellow arrows indicate muscle cells, and white triangles denote fibroblasts. White dashed lines outline the fin shape, while yellow dashed lines mark the amputation sites. Insets display higher magnification views. (C) Immunostaining of MuSC-derived cell identities in (B) using PRRX1 (fibroblast marker), MHC (muscle marker), and SOX9 (chondrocyte marker). Triangles indicate fibroblasts, open triangles denote chondrocytes, and arrows point to muscle cells. Scale bars: 1 mm (B); 50 μm (C).

We previously showed that TGF-β signaling modulates MuSCs fate during axolotl tail regeneration, where activation via caTGFBR1 biases MuSCs toward fibroblast and chondrocyte lineages (Figure 6Aa), and inhibition via SMAD7 favors myogenic differentiation (Figure 6Ac). Notably, the phenotypes observed upon TGF-β manipulation mirror those resulting from ERK perturbation (Figures 6Ab and 6Ad). Given that the upstream hierarchy between ERK and TGF-β is tissue specific,^38,39,40,41,42^ we next first investigated whether TGF-β regulates ERK activation in MuSCs. Using a double-transgenic line (*Pax7:CreERT2/CAGGS:loxP-STOP-loxP-caTgfbr1-Cherry*) to specifically activate TGF-β signaling in MuSCs (Figures S10A and S10B), we observed that caTGFBR1 induction had no influence on ERK activity in *Pax7*+ MuSCs compared to controls (Figures S10Cc, S10Ca, S10D). In contrast, expression of caMEK robustly activated ERK signaling in the positive control (Figure S10Cb). These findings suggest that, in MuSCs, the TGF-β signaling pathway dose not activate ERK.

To test whether ERK functions upstream of TGF-β to regulate MuSC plasticity, we employed two triple-transgenic lines tTG-Pax7 (Pax7:CreERT2/CAGGS:loxP-BFPnls-STOP-loxP-GFP-caMek-Cherry/CAGGS:GFP -Smad2 and Pax7:CreERT2/CAGGS:loxP-BFPnls-STOP-loxP-GFP-caMek-Cherry/CAGGS:GFP- Smad3) to assess the impact of ERK activation on TGF-β downstream targets localization of SMAD2 and SMAD3 (Figure S10E and S10F). We found that ERK activation enhanced the phosphorylation and nuclear translocation of both SMAD2 and SMAD3 (Figures S10G-S10I), suggesting that ERK promotes TGF-β signaling by facilitating SMAD2/3 activation.

Finally, to validate that ERK modulates MuSC plasticity through the TGF-β pathway in vivo (Figure 6B), we generated triple-transgenic animals (*Pax7:CreERT2/CAGGS:loxP-STOP-loxP-caTgfbr1-Cherry/CAGGS:loxP-STOP-lox P-dnMek-GFP*) to determine if TGF-β activation could rescue the phenotype induced by ERK inhibition (Figure 6C and 6D). 14 days post-4-OHT treatment, co-localization of dnMEK and caTGFBR1 was confirmed in *Pax7*+ MuSCs (Figures S11A and S11B). Analysis of GFP+/Cherry+ double-positive cells at 21 dpa revealed cells with chondrocyte-like and fibroblast-like morphologies (Figures 6Ea and 6Eb), which immunostaining confirmed as SOX9+ chondrocytes and PRRX1+ fibroblasts (Figures 6Fa and 6Fb). These findings show that TGF-β activation rescues the phenotype induced by ERK inhibition.

**Figure 6.**
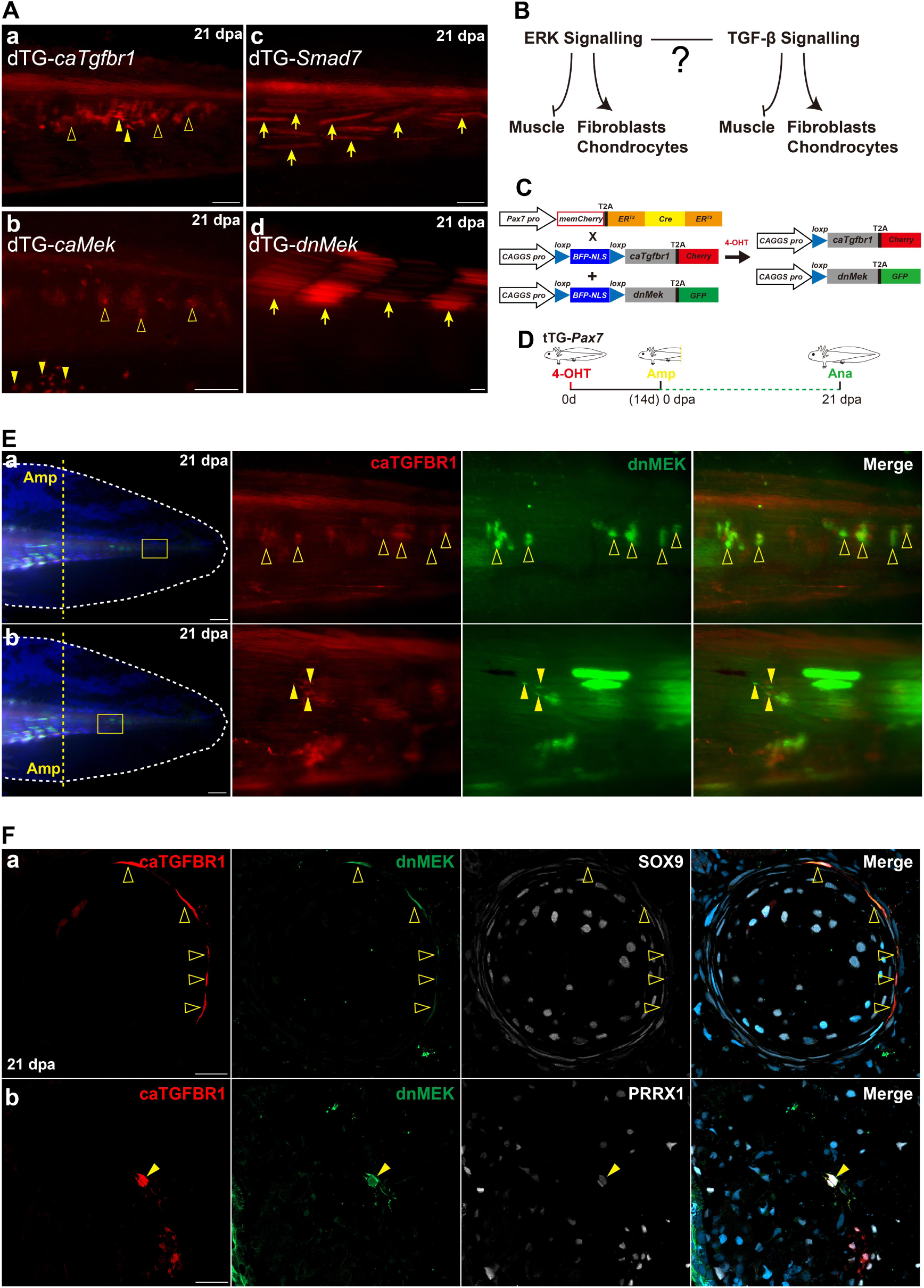
TGF-β activation rescues the phenotype induced by ERK inhibition. (A) Representative images showing both sustained TGF-β activation (a) (dTG-*caTgfbr1*, n=11) or ERK activation (b) (dTG-*caMek*, n=4) drove MuSC plasticity towards fibroblasts and chondrocytes, and both sustained TGF-β inhibition(c) (dTG-*Smad7*, n=8) or ERK inhibition (d) (dTG-*dnMek*, n=3) induces muscle formation at 21 dpa. (B) Schematic depicting the proposed regulatory relationship between ERK and TGF-β signaling. (C, D) Schematic of the experimental strategy combining ERK inhibition with TGF-β activation in the *Pax7*+ MuSC lineage during tail regeneration. (E) Representative images of regenerated *Pax7*+ MuSC-derived cells under combined ERK inhibition and TGF-β activation (n=5). Yellow dashed lines mark the amputation sites, and white dashed lines delineate the fin shape; insets show enlarged views. (F) Immunostaining characterization of cell identity in double-positive cells from (E). Scale bars: 250 μm (A); 1 mm (E); 50 μm (F). Empty triangles indicate chondrocytes; filled triangles indicate fibroblasts; arrows indicate muscle cells.

Collectively, our in vitro and in vivo data support a model that ERK regulates MuSC plasticity during axolotl fin regeneration through modulation of TGF-β signaling.

### SMAD2, not SMAD3 mediates ERK–TGF-β regulation of MuSC plasticity towards a fibroblast fate

TGF-β signaling is typically transduced by the effectors SMAD2 and SMAD3, which have been shown to play distinct roles in different tissues. For instance, SMAD2, but not SMAD3 is required to mediate TGF-β signaling during axolotl limb regeneration.^43^ TGF-β on macrophage function involves SMAD3, not SMAD2.^44^ Hence, to investigate the specific roles of SMAD2 and SMAD3 in ERK-regulated MuSC plasticity during axolotl tail regeneration, we constructed the transgenic line *CAGGS:GFP-Smad2/3* to monitor SMADs activity in vivo via nuclear GFP localization (Figure S12A). Immunostaining of uninjured axolotl tail muscle revealed that SMAD3, but not SMAD2, accumulated in the nuclei of MHC+ cells (Figures S12B-S12C), whereas both SMAD2 and SMAD3 showed low activity in *Pax7*+ MuSCs (Figures S12D-S12E). These observations suggest that, during muscle cell generation, TGF-β predominantly regulates SMAD3. Based on these findings, we hypothesized that SMAD2, rather than SMAD3, is involved in mediating the ERK-TGF-β regulation of MuSC differentiation towards the fibroblast-chondrocyte lineage.

To test this hypothesis, we generated constitutively active forms of axolotl SMAD2 and SMAD3 that mimic phosphorylation-induced nuclear translocation. We introduced a serine-to-tyrosine substitution at position 306 in SMAD2 (caSMAD2) and at position 270 in SMAD3 (caSMAD3) to disrupt the auto-inhibitory domain and facilitate nuclear localization.^45^ Both caSMAD2 (S306Y) and caSMAD3 (S270Y) mutants promoted robust nuclear translocation (Figures S13A-S13D). We then constructed triple-transgenic animals (*Pax7:CreERT2/CAGGS:loxP-STOP-loxP-GFP-caSmad2/CAGGS:loxP-STOP-loxP- Smad7-Cherry* and *Pax7:CreERT2/CAGGS:loxP-STOP-loxP-GFP-caSmad3/CAGGS:loxP-STOP-loxP-S mad7-Cherry*) to assess whether the activated SMAD mutants could override the inhibitory effect of SMAD7 on endogenous SMAD2/3 signaling and thereby regulate MuSC plasticity (Figure 7A-7C). SMAD7 overexpression inhibited endogenous SMAD2 and SMAD3, allowing us to isolate the effect of each activated mutant. Fourteen days post-4-OHT treatment, co-localization of Cherry (SMAD7) and GFP (SMAD2/3) was detected in axolotl tail muscle compartments (Figures S14A), and immunostaining confirmed their co-expression in *Pax7*+ MuSCs (Figures S14B).

**Figure 7.**
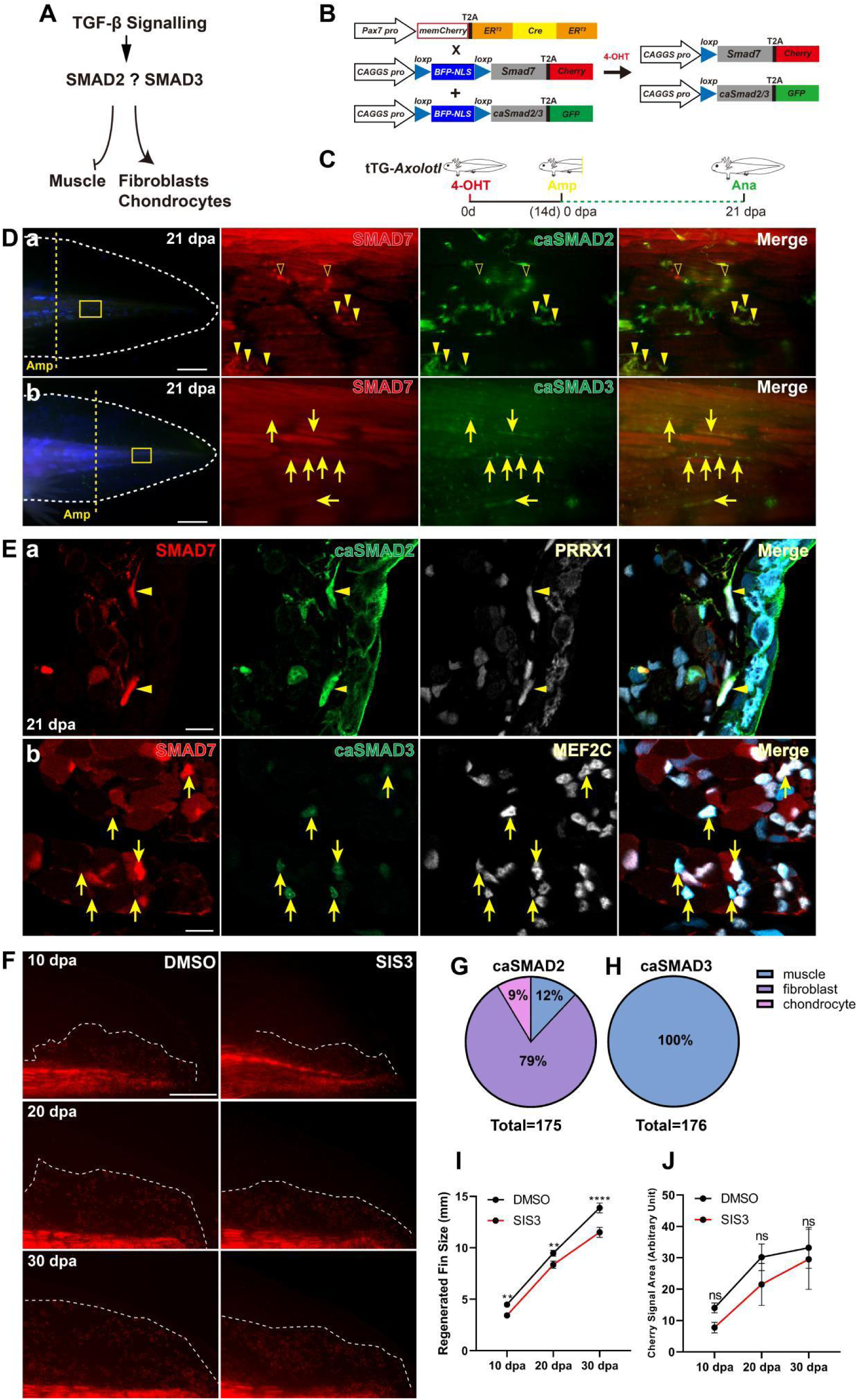
Continuous activation of SMAD2, not SMAD3 rescues TGF-β inhibition regulated phenotype. (A) Schematic depicting the proposed TGF-β signaling mediated SMAD2/SMAD3 on cell fate decision. (B, C) Schematic of the experimental strategy combining inhibiting endogenous TGF-β activity and continuously activate SMAD2 or SMAD3 in the *Pax7*+ MuSC lineage during tail regeneration. (D) Live imaging showing caSMAD2 (a) (n=6), not caSMAD3 (b) (n=6) regulates *Pax7*+ MuSCs plasticity towards fibroblasts during tail regeneration. Triangles indicate fibroblasts; arrows indicate muscles. Yellow dashed lines indicate amputation sites; white dashed lines indicate fin shape; enlarged images were shown at right panels. (E) Immunostaining characterization of cell identity in (D). PRRX1 marks fibroblasts; MEF2C marks myoblasts and muscle nucleus. Triangles indicate SMAD7 and caSMAD2 co-expressed cells; arrows indicate SMAD7 and caSMAD3 co-expressed cells. (F) Live imaging showing fin fibroblasts generation during regeneration under SMAD3 inhibition (SIS3, n=6). Dashed lines indicate fin regions. (G, H) Quantitative analysis of cell type percentage in (Da) caSMAD2+/SMAD7+ and (Db) caSMAD3+/SMAD7+ cells. (I, J) Quantitative analysis of regenerated fin size and Cherry signal areas in (F) (mean ± SEM; each dot represents a cell; **p < 0.01; ****p < 0.0001; ns, not significant). Scale bars: 2 mm (D); 20 μm (E); 1 mm (F).

At 21 days post-amputation, lineage tracing of GFP+/Cherry+ double-positive cells revealed distinct cell fate outcomes depending on the activated SMAD mutant. In tails expressing SMAD7 together with constitutively active SMAD2 (caSMAD2), 79% of the traced cells exhibited a typical fibroblast-like morphology and co-localized with the fibroblast marker PRRX1 (Figure 7Da, 7Ea, 7G). A minority of the cells adopted alternative fates, with 9% displaying chondrocyte-like morphology and 12% exhibiting muscle-like characteristics. In contrast, when SMAD7 was co-expressed with constitutively active SMAD3 (caSMAD3), 100% of the traced cells were integrated into muscle fibers and co-localized with the muscle-specific marker MEF2C (Figure 7Db, 7Eb, 7H). Furthermore, chemical inhibition of endogenous SMAD3 in *Pax7*+ MuSCs using SIS3 (at a dosage previously established in axolotls)^43,46^ resulted in a slight reduction in tail regeneration length but did not significantly affect the generation of MuSC-derived fibroblasts (Figure 7F, Figures 7I and 7J).

Together, our in vivo genetic manipulation and pharmacological inhibition experiments indicate that SMAD2, rather than SMAD3, is the critical downstream effector mediating ERK-TGF-β regulation of MuSC plasticity toward a fibroblast fate during axolotl tail regeneration.

## DISCUSSION

### Phased ERK function coordinates tail regeneration

In MuSCs, the ERK signaling pathway has been identified as a tissue damage-induced stress signal that regulates their the transition of from a quiescent to an active state.^47,48,49^ Research has shown that ERK signaling promotes the proliferation of myoblasts while inhibiting their differentiation into muscle cells.^50,51,52,53^ Nonetheless, no studies have reported ERK’s role in directing the differentiation of MuSCs toward non-muscle lineages. ERK dynamics has been shown to play different effects even within the same tissue at different developmental stages. For instance, during zebrafish periderm morphogenesis, ERK first regulates proliferation and then cell size to coordinate embryo elongation.^54^ Except for development, our study revealed a phased ERK regulatory mechanism directing MuSCs plasticity towards fibroblast lineages which coordinates axolotl tail regeneration.

Sustained ERK activation is indispensable in other regenerative contexts. For instance, upon tissue loss, spiny mice (Acomys) ears can regenerate with no scar tissue, during which, a robust high level of ERK is initiated and maintained till new tissue formation.^55^ In newt myotube cell lines, sustained ERK activation is essential for cell cycle re-entry and epigenetic modifications, a phenomenon not observed in mammalian counterparts.^56^ Similarly, in American cockroach limb regeneration, sustained ERK activation is necessary for blastema formation and progenitor cell differentiation, and its disruption impedes further regeneration.^57,58^ FGF signaling is indicated to be the upstream regulator of ERK, since FGF signaling can regulate stem cell fate through ERK singnaling,^59^ and inhibition of FGF pathway hinders early regeneration events, results in no blastema formation.^60,61^ Although sustained ERK activity and its upstream regulators have been studied, the essential role of ERK activation on MuSCs fate and the importance of its inhibition for coordinating regeneration was not examined. We observed that injury-induced ERK activation is sustained turned till 6 dpa (Figure 1F) during the regeneration of axolotl tails. During this time, ERK directs the differentiation of MuSCs towards trunk fibroblasts. From 10 dpa on, ERK activity is inhibited in MuSCs and their cell progeny, and this inhibition is required to generate fin fibroblasts. Our study provided evidence that both sustained ERK activation and subsequent inhibition are indispensable for coordinated trunk and fin fibroblast generation during axolotl tail regeneration.

### Muscle stem cell plasticity regulation

ERK activation is known to occur rapidly following various tissue injuries, including nerve, epidermis, blood vessel,^62,63,64^ and is important for neural specification and embryonic stem cell lineage commitment.^65,66^ Although ERK is reported to activate MuSCs from a quiescent to proliferative state,^47^ whether it’s involved in stem cell plasticity transduction is unknown. In our study, we discovered a new function of ERK on MuSCs lineage commitment to fibroblast lineage.

To target ERK-regulated muscle stem cell plasticity, it is essential to understand the underlying mechanisms. First, for a stem cell to switch to a different identity, it must first erase its original markers. In our study, we observed that MuSCs lost their PAX7 identity as soon as pERK levels were elevated upon injury. Overexpression experiments revealed that ERK activation represses PAX7 expression (Figure S6B-S6G), this observation is also supported by an in vitro study.^34^ Knockout PAX7 is sufficient to induce MuSCs transit to trunk fibroblasts. In addition to PAX7, evidence suggests that sustained ERK activation downregulates the muscle-specific gene *Sox6* in a different salamander species.^56^ Beyond MuSCs, we also observed pan-upregulation of ERK activity during axolotl tail regeneration, including in neural progenitor cells (data not shown). Thus, future work should investigate whether this mechanism is conserved across different cell types and explore the molecular pathways involved in identity reprogramming.

Second, the acquisition of a new cell identity requires guidance from specific signaling pathways that activate a distinct combination of genes defining the new cell type. Our data demonstrate that the ERK-regulated TGF-β/SMAD2 pathway is a key molecular switch driving MuSC plasticity toward trunk fibroblast lineage during axolotl tail regeneration (Figures 6-7). However, as a downstream target of ERK, TGF-β does not regulate MuSC plasticity under homeostatic conditions (Figure S7D-S57F). This suggests that additional ERK-specific downstream targets are involved in regulating plasticity during the injury response. For instance, CK-2.^57^ Future studies also need to identify the potential mechanisms that mediate MuSC plasticity, such as epigenetic modifications^56,67,68^ and specific transcription factors.^69,70^ Then, ERK inhibition is required for trunk fibroblasts to migrate into fins. We found PRRX1 is highly expressed in fin fibroblasts, and ERK activation inhibits PRRX1 expression. So, ERK inhibition facilitates fin fibroblasts formation is possibly through PRRX1. Future study to test the function of PRRX1 on trunk fibroblasts transit to fin is needed.

## RESOURCE AVAILABILITY

### Lead contact

Requests for further information and resources may be directed to and will be fulfilled by the lead contact, Jifeng Fei.

### Materials availability

Plasmids generated in this study will be distributed on request.

### Data and code availability

• This paper does not report original code.
• Single cell sequencing data can be accessed from the National Genomics Data Center’s Genome Sequence Archive (https://ngdc.cncb.ac.cn/gsa, accession CRA019184 for Smart-seq2 datasets).
• Any additional information required reported in this paper is available from the lead contact upon request.

## ACKNOWLEDGEMENTS

We thank the animal facility staff, Guoqing Liu and Zhenyao Wu, for their outstanding technical support. We also acknowledge the expertise and assistance provided by the central equipment platform at Guangdong Provincial People’s Hospital and the computing resources supplied by China National GeneBank. Our gratitude extends to Dr. Christopher L. Antos for his insightful scientific discussions and valuable comments on the manuscript.

## AUTHOR CONTRIBUTIONS

Conceptualization: CY, J-FF

Methodology: CY, XL, J-FF

Investigation: CY, XL, LW, LS

Visualization: CY, XL, J-FF

Funding acquisition: CY, J-FF

Project administration: CY, J-FF

Supervision: J-FF

Writing—original draft: CY

Writing—review & editing: CY, J-FF, AC, XL

## STAR*Methods

### Key resources table

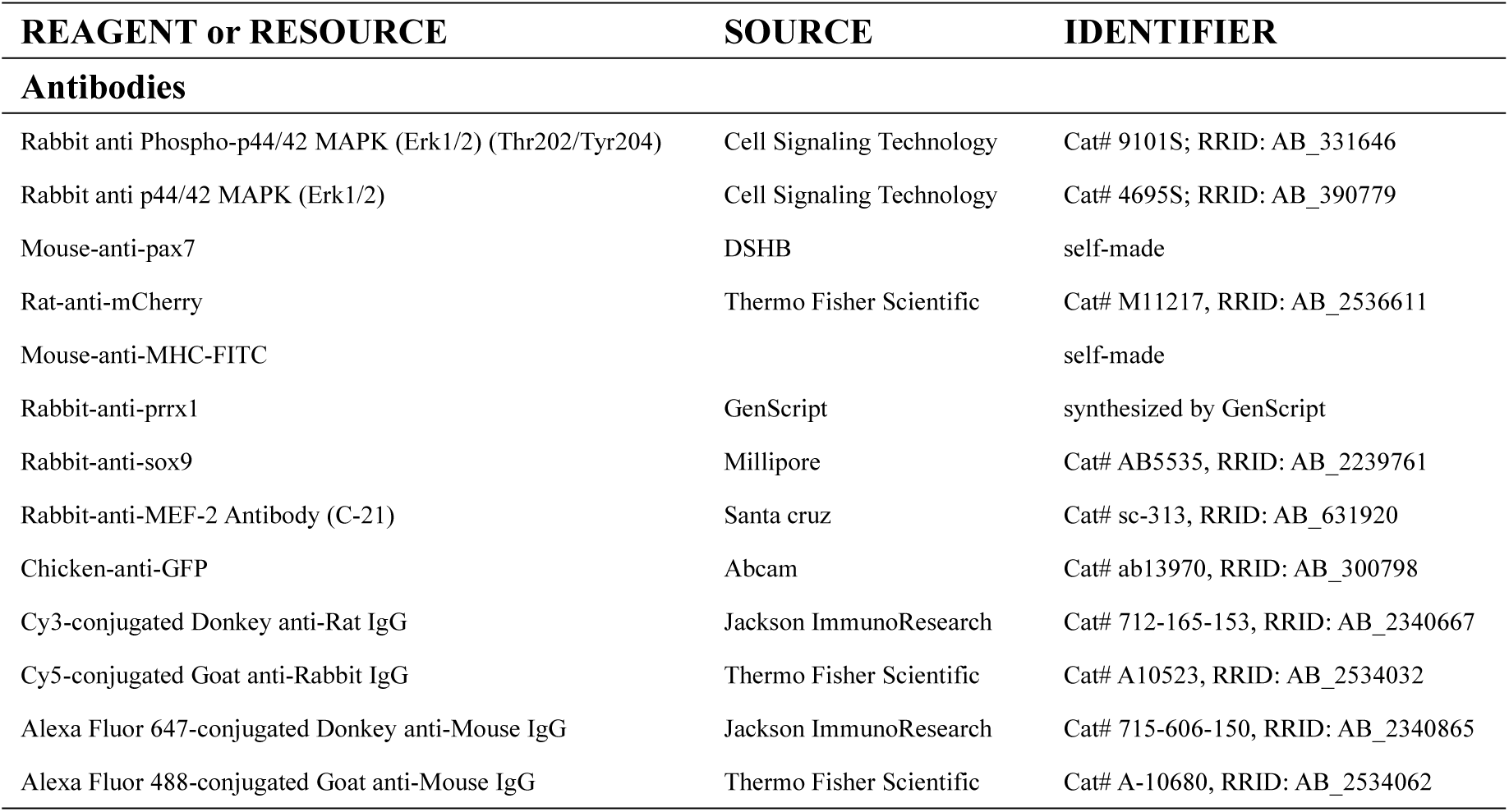
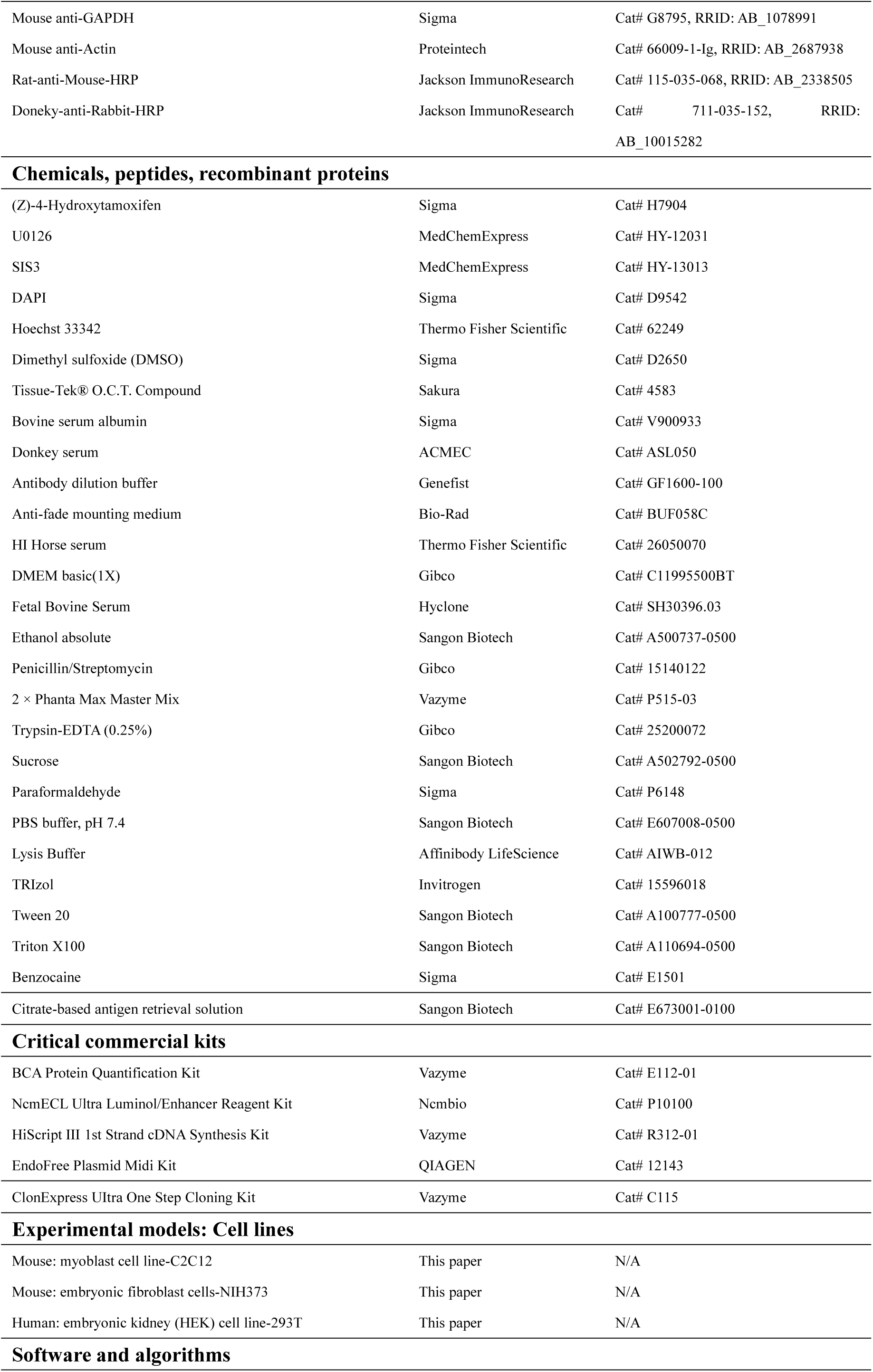
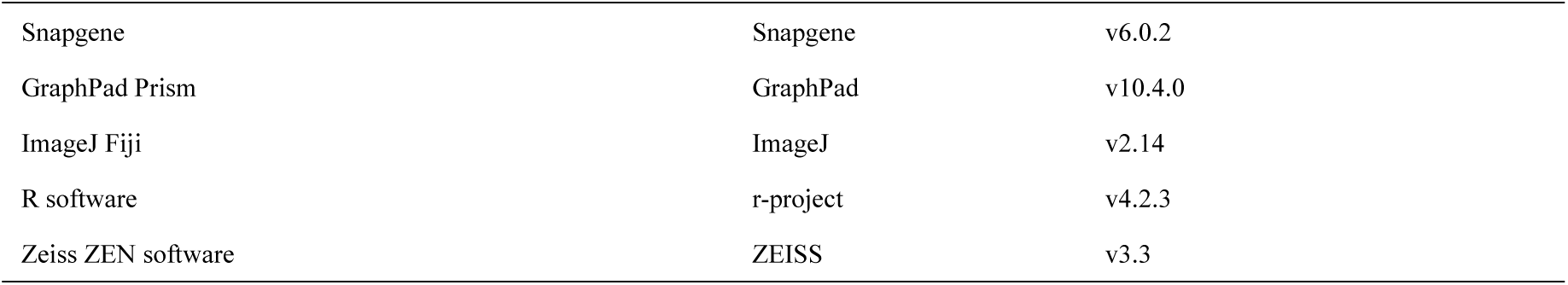

## METHOD DETAILS

### Animals

In this study, wild type (*d/d*) axolotl (*Ambystoma mexicanum*) strain along with two previously described transgenic lines *Pax7:CreER^T2^* (*Pax7:Pax7*-ORF^b^Δ-*P2A*-*memCherry*-*T2A*-*ER^T2^-Cre-ER^T2^*) and *CAGGS:reporter* (*CAGGS:LoxP-GFP-STOP-LoxP-Cherry*)^21,71^ were obtained from Elly M. Tanaka’s laboratory at the Research Institute of Molecular Pathology in Vienna, Austria. The *Pax7:Cherry* knock-in/knockout axolotls, *CAGGS:loxp-BFP-STOP-loxp-caTgfbr1-T2A-Cherry* and *CAGGS:loxp-BFP-STOP-loxp-Smad7-T2A-Cherry* were constructed in our lab’s previous work.^13,36^ Additional transgenic strains were generated in this work according to previous published protocols.^71^ The axolotls were maintained at 20°C in freshly dechlorinated tap water and were fed daily. All experimental procedures were performed in accordance with the relevant Chinese animal welfare regulations and approved by the Biomedical Research Ethics Committee of Guangdong Provincial People’s Hospital (approval number KY2024-192-01).

### Cloning

#### Tol2-CAGGS:loxp-BFP-STOP-loxp-caMek/dnMek/caCtnnb1-T2A-Cherry-PA, Tol2-CAGGS:loxp-STOP-loxp-psMek-T2A-Cherry-PA, ***Tol2-CAGGS:loxp-BFP-STOP-loxp-caSMAD2/3-T2A-GFP-PA*:**

CDS of *Mek1, Smad2, Smad3* were PCR-amplified from axolotl fin cDNA libraries using *Mek1, Smad2, Smad3* primer pairs, *caCtnnb1* was amplified according to published study^32^ using *caCtnnb1* primer pair. *psMek* sequence was synthesized by GENEWIZ company. *Mek1* was mutated to either *caMEK* or *dnMEK* by GENEWIZ company. Next, these genes were used as templates were amplified by PCR to add homologous arms with indicated primers listed in Supplementary table 1, They were then inserted into the vector *Tol2-CAGGS:loxp-BFP-nls-STOP-loxp-T2A-Cherry/GFP-PA* or *Tol2-CAGGS:loxp-STOP-loxp-T2A-Cherry-PA*^13^ using the in-fusion cloning method to generate *Tol2-CAGGS:loxp-BFP-nls-STOP-loxp-caMEK/dnMek/caCtnnb1-T2A-Cherry-PA* and *Tol2-CAGGS:loxp-STOP-loxp-psMEK-T2A-Cherry-PA and Tol2-CAGGS:loxp-BFP-nls-STOP-loxp-Smad2/3-T2A-GFP-PA*. Plasmids *Tol2-CAGGS:loxp-BFP-nls-STOP-loxp-caSmad2/3-T2A-GFP-PA* were further constructed by PCR amplification with the corresponding mutation primer pairs (*caSmad2, caSmad3*), using *Tol2-CAGGS:loxp-BFP-nls-STOP-loxp-Smad2/3-T2A-GFP-PA* as template with a high fidelity DNA polymerase.

#### Tol2-CAGGS:GFP-Smad2/Smad3-PA

Vector *Tol2-CAGGS:GFP-PA* was digested with NheI and BamHI. *GFP* and *Smad2/3* genes were used as templates and amplified using primer pairs HR-*GFP-Smad2/3*. They were then inserted into the digested vector using the in-fusion cloning method to generate *Tol2-CAGGS:GFP-Smad2/Smad3-PA*.

All of the final constructs were confirmed by Sanger sequencing. The primer sequences for cloning are listed in the Supplementary table 1.

#### Cell culture

C2C12, NIH3T3 and HEK 293T cell lines were cultured at 37℃, 5% CO_2_ and 95% humidity in DMEM medium supplemented with 10% FBS and 1% penicillin-streptomycin. Cells were split to 30% density using 0.25% trypsin and transfected with indicated plasmids using lipofectamine (Thermofisher L3000-015) at 70% confluency following the manufacturer’s guidance. The expression of the transfected constructs was evaluated by fluorescence imaging. C2C12 myoblasts were induced to differentiate into myotubes using differentiation medium, which consisted of DMEM supplemented with 2% horse serum.

#### Drug treatment

A 20 mM stock solution of 4-OHT was prepared by dissolving the compound in DMSO, aliquoted, and stored at -80°C. For treatment, the 4-OHT stock was diluted to 2 µM in water. Drug administration was carried out over three consecutive days, interspersed with one-day intervals, and treatment efficacy was assessed via fluorescence microscopy four days after the final treatment. U0126-EtOH and SIS3 were each dissolved in DMSO to yield stock solutions at 30 mM and 20 mM, respectively. For U0126 treatment, 2-3 cm size animals were incubated in a 40 µM U0126 solution at 6 days post-amputation, following the concentration reported by Hurtado et al.^72^ For SIS3 treatment, a preliminary dose-response assessment was performed using concentrations of 5 µM, 4 µM, and 3 µM according to the literature.^43^ Treatments with 5 µM resulted in 100% mortality within 24 hours, while those with 4 µM caused significant morbidity, as evidenced by bubble formation and abnormal buoyancy. Animals treated with 3 µM displayed minimal adverse effects; thus, this concentration was selected for subsequent experiments. After tail amputation, 2-3 cm animals were immersed in 3 µM SIS3, while corresponding control groups received an equivalent volume of DMSO. All drug solutions were refreshed daily, and fins were monitored at designated time points using a fluorescent stereomicroscope.

#### RNA extraction, cDNA synthesis and Real-time quantitative PCR

Total RNA was extracted by TRIzol methods. Subsequently, 1 μg of RNA was wiped genomic DNA and reverse-transcribed with HiScript III 1st Strand cDNA Synthesis Kit. RT-qPCR was performed and normalized using the internal control for GAPDH. The relative expression levels of target genes were calculated using the 2^-ΔΔCt^ method. The primer sequences used were as follows: GAPDH: 5′- GACAAGGCATCTGCTCACCT-3′ (forward) and 5′- ATGTTCTGGTTGGCACCTCT-3′ (reverse); Pax7: 5′- TACACCCGTGAAGAGTTGGC-3′ (forward) and 5′- TTCCTGTGGGTGGAAATCCG-3′ (reverse).

#### Western blotting

Protein lysates were prepared from axolotl tissues treated with either DMSO or U0126 by homogenization in a commercial lysis buffer. The samples underwent sonication in 3-second pulses with 3-second pauses, repeated for a total of 1 minute at 40 W (xmcsbyq, XM-650DT). Following sonication, the lysates were centrifuged at 12,000 rpm for 20 minutes at 4℃, and the supernatants were collected. Protein concentrations were determined using the BCA assay according to the manufacturer’s instructions. For electrophoretic analysis, 30 μg of protein per sample was separated on a 4-12% SDS-PAGE gel (ACE, F15412MGel) by running at 160 V for 60 minutes. Proteins were then transferred onto a PVDF membrane (Immobilon, IPVH00010) using a semi-dry transfer system set to 20 V for 15 minutes. Membranes were blocked with 5% BSA at room temperature for 1 hour. The membranes were then incubated overnight at 4°C with primary antibodies: rabbit anti-pERK, rabbit anti-ERK, and mouse anti-β-actin. After thorough washing, membranes were incubated at room temperature for 1 hour with secondary antibodies: rat anti-mouse-HRP or donkey anti-rabbit-HRP. Following three washes with TBST (0.1% Tween), immunoreactive bands were visualized using the NcmECL Ultra Luminol/Enhancer Reagent kit and imaged with the Touch Imager (E-BLOT).

#### Immunofluorescence

Air-dried cryosections or fixed cells were initially rinsed with phosphate buffered saline containing 0.3% Triton X-100 (PBST). For samples requiring antigen retrieval, sections were incubated at 85°C for 10 minutes in a citrate-based antigen retrieval solution. Following retrieval, the slides were washed with PBST to eliminate any residual retrieval buffer. Subsequently, sections were blocked with 5% donkey serum to minimize nonspecific binding. Primary antibody incubation was then performed overnight at 4°C, with antibodies diluted in an optimized antibody dilution buffer. After rinsing with PBST, samples were incubated at room temperature for 2 hours with the corresponding secondary antibodies and DAPI. The primary and the secondary antibodies used were listed above. Finally, the sections were mounted using an anti-fade mounting medium prior to imaging.

#### Optogenetic controlling of endogenous ERK activity

The Optogenetic tool psMEK used in this study was synthesized according to published work.^37,73^ When psMEK was illuminated with a wavelength of 488 nm, dimerized photoswitchable protein Dronpa (green fluorescent) was dissociated and exposed the caMEK site. Conversely, when psMEK was illuminated with a wavelength of 405 nm, Dronpa protein would re-dimerize and blocked the activity of caMEK. The animals were anesthetized in 0.03% (w/v) benzocaine and sited on a wet petri-dish prior to illumination. The animals were then illuminated at either 405 nm or 488 nm for 30 minutes using an illuminator (FluoCa, SN B36638). Control animals were kept in dark environment.

#### scRNA-seq analysis

Single-cell RNA sequencing (scRNA-seq) data were retrieved from the National Genomics Data Center’s Genome Sequence Archive (https://ngdc.cncb.ac.cn/gsa, accession CRA019184 for Smart-seq2 datasets). Pathway activation scores were computed according to methodologies previously described^13^.

#### Imaging

Axolotl images were obtained using bright-field and fluorescence modalities on an Olympus stereomicroscope (SXZ10, SXZ16) equipped with a FLuoCa fluorescent illuminator. For visualizing fluorescent signals in sections and cultured cells, a confocal microscope (LSM980) fitted with either 10 × or 20 × objectives was employed.

#### Statistical analysis

Data were analyzed using GraphPad Prism software, with all values reported as the mean ± standard error of the mean (SEM). For pairwise comparisons of independent groups, a two-tailed unpaired Student’s *t*-test was employed. When comparing more than two groups, a one-way ANOVA was conducted, followed by either Dunnett’s test or Fisher’s Least Significant Difference (LSD) post hoc analysis as appropriate. A *P*-value below 0.05 was deemed statistically significant.

